# Hysteresis in PIF4 and ELF3 dynamics dominates warm daytime memory in Arabidopsis

**DOI:** 10.1101/2021.10.19.465000

**Authors:** Germán Murcia, Cristina Nieto, Romina Sellaro, Salomé Prat, Jorge J. Casal

## Abstract

Plants may experience large diurnal temperature fluctuations. Our knowledge of the molecular mechanisms of integration of these fluctuations and the resulting growth patterns is limited. Here we show that hypocotyl growth during the night responded not only to the current temperature but also to preceding daytime temperatures, revealing a memory of previous conditions. Daytime temperature affected the nuclear levels of PHYTOCHROME INTERACTING FACTOR 4 (PIF4) and LONG HYPOCOTYL 5 (HY5) during the next night. These jointly accounted for the observed growth kinetics, whereas memory of prior daytime temperature was impaired in the *pif4* and *hy5* mutants. *PIF4* promoter activity largely accounted for the temperature dependent changes in PIF4 protein levels. Noteworthy, the decrease in *PIF4* promoter activity triggered by cooling required a stronger temperature shift than the increase caused by warming. This hysteretic pattern required EARLY-FLOWERING 3 (ELF3). Warm temperatures promoted the formation of nuclear condensates of ELF3 in hypocotyl cells during the afternoon but not in the morning. These nuclear speckles showed poor sensitivity to subsequent cooling. We conclude that ELF3 achieves hysteresis and drives the *PIF4* promoter into the same behaviour, enabling a memory of daytime temperature conditions.

## Introduction

Warmer temperatures within the physiological range can selectively increase or decrease the growth of different organs, leading to modifications in plant architecture or thermomorphogenesis (Quint et al., 2016; Casal and Balasubramanian, 2019). Thermomorphogenesis occurs in crop species, highlighting the need to understand this process in further deepness in the current context of global warming (Casal and Balasubramanian, 2019). A well-established model in thermomorphogenesis, useful to understand the basic mechanisms, is the enhanced growth of the hypocotyl in *Arabidopsis thaliana* seedlings in response to non-stressful warm temperatures (Gray et al., 1998).

The photo-sensory receptor phytochrome B (phyB) (Jung et al., 2016; Legris et al., 2016), the clock protein EARLY FLOWERING 3 (ELF3) (Jung et al., 2020) and the transcription factor PHYTOCHROME INTERACTING FACTOR 7 (PIF7) (Chung et al., 2020) are the three plant temperature sensors involved in the control of hypocotyl growth identified so far. Warm temperatures accelerate the rate of thermal reversion of active phyB to its inactive conformer (Jung et al., 2016; Legris et al., 2016; Burgie et al., 2021). ELF3 is a component of the evening complex and warm temperatures reduce its binding to the target gene promoters (Box et al., 2015; Ezer et al., 2017; Silva et al., 2020). ELF3 contains a predicted prion domain with a polyglutamine repeat, which is important for the reversible phase transition from active to the inactive state of ELF3 under warm conditions (Jung et al., 2020). According to Jung et al. (2020), warm temperatures can enhance the formation of nuclear speckles containing ELF3, but Ronald et al. (2021) reported the opposite pattern. Therefore, there is controversy about the link between these a sub-nuclear bodies and ELF3 activity. Warmth-induced changes in the RNA hairpin present at the 5′-untranslated region of the *PIF7* transcript favour its translation, increasing PIF7 protein abundance (Chung et al., 2020). phyB and ELF3 signalling converge on PIF4, as phyB physically interacts and negatively regulates PIF4 protein stability (Cheng et al., 2021) and the evening complex binds the *PIF4* gene promoter reducing its expression during early night (Nusinow et al., 2011). Therefore, both *PIF4* expression (Koini et al., 2009; Stavang et al., 2009; Box et al., 2015) and PIF4 protein stability (Foreman et al., 2011) increase under elevated temperatures. In addition, ELF3 sequestrates PIF4 by direct physical interaction preventing PIF4 binding to its transcriptional targets (Nieto et al., 2015).

CONSTITUTIVELY PHOTOMORPHOGENIC 1 (COP1), ELONGATED HYPOCOTYL 5 (HY5) and LONG HYPOCOTYL IN FAR-RED (HFR1) are among the several regulators of PIF4 activity. COP1 is a RING E3 ligase that is required for the hypocotyl growth response to warm temperatures (Delker et al., 2014). Warm temperature increases nuclear accumulation of COP1 (Park et al., 2017) and this enhances the expression of *PIF4* (Gangappa and Kumar, 2017). HY5 binds the *PIF4* promoter to negatively regulate its activity (Delker et al., 2014). HY5 also competes with PIF4 in its binding to the target gene promoters (Toledo-Ortiz et al., 2014), while warm temperatures inhibit the expression of the *HY5* gene and can lower the HY5 protein stability (Catalá et al., 2011; Delker et al., 2014; Toledo-Ortiz et al., 2014). HFR1 is stabilised under warm temperatures (Romero-Montepaone et al., 2021) and inhibits PIF4 and PIF7 binding to its targets, by direct physical interaction (Hornitschek et al., 2009; Sandi Paulišić et al., 2021). The transcription factor PIF4 (Crawford et al., 2012; Sun et al., 2012) and PIF7 (Fiorucci et al., 2020) transcription factors bind auxin synthesis gene promoters to increase auxin levels in the warmth. Auxin produced in the cotyledons travels down to the hypocotyl to promote growth (Bellstaedt et al., 2019).

In nature, temperatures typically fluctuate between day and night. We have recently observed that night temperature information stored in phyB affects hypocotyl growth during the subsequent photoperiod (Murcia et al., 2020). However, we ignore whether the reciprocal control is also true; i.e. whether temperature responsive hypocotyl elongation at night depends on the temperature experienced during the preceding photoperiod. Here we report that nighttime hypocotyl growth and gene expression depend not only on the temperature during the night itself but also on former daytime temperature.

## Results

### Nighttime growth depends on daytime temperatures

To investigate whether the growth of the hypocotyl during the night responds exclusively to the current temperature or is also affected by the conditions experienced during the previous photoperiod, plants of the Col wild type (WT) were exposed to all possible combinations between three daytime temperatures and three nighttime temperatures (10, 20 and 28°C). Furthermore, to analyse whether the status of photo-sensory receptors modify the influence of previous temperature, plants were exposed to simulated sunlight or shade conditions during daytime and either a far-red light pulse (EOD FR, 10 min) or no light pulse at dusk. As expected, warmer nights accelerated nighttime growth (Fig. 1A, Table S1A). More importantly, for all nighttime conditions, warmer daytime temperatures caused faster nighttime growth (Fig. 1 A, Table S1A). EOD FR or daytime shade accelerated hypocotyl growth during the night (Table S1A). Significant interactions indicate that the effect of daytime temperature was stronger when nighttime temperatures were warmer and in seedlings exposed to EOD FR (Table S1A). Taken together, these results indicate that nighttime growth depends not only on the nighttime temperature itself but also on prior daytime temperature, particularly under the conditions that elicit faster nighttime growth (warm night, daytime shade, EOD FR). Based on these observations, in subsequent experiments we used daytime shade and EOD FR to optimise the analysis. This phenomenon should not be confused with the known effects of day / night temperature differentials, which actually impact on daytime growth (Bours et al., 2013, 2015).

**Figure 1.**
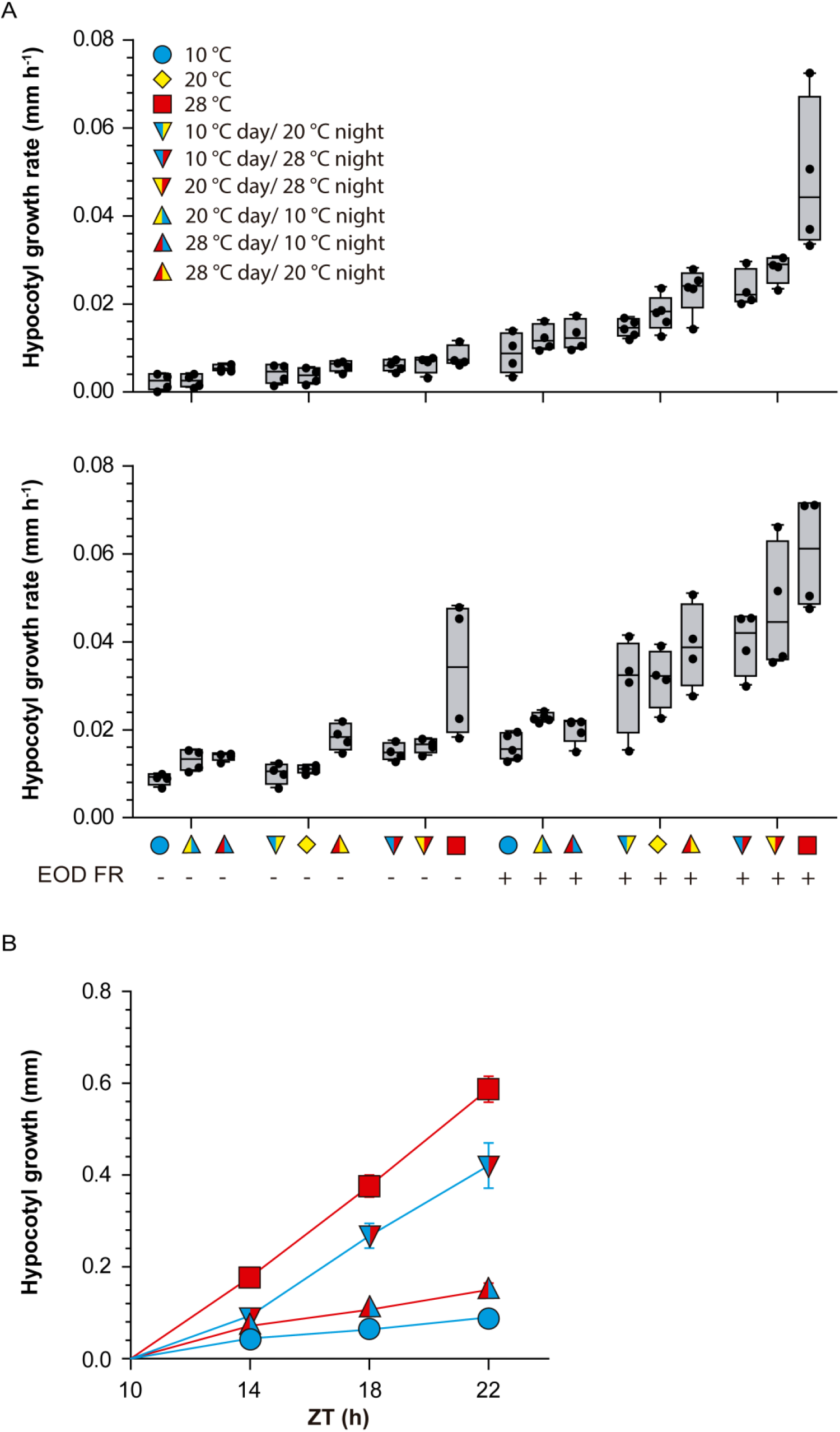
Daytime temperatures affect nighttime hypocotyl growth. **(A)** Hypocotyl growth rate measured in Col-0 seedlings during the night (ZT 10h to ZT 24h). The seedlings received all possible combinations of temperature during the preceding photoperiod (10, 20 or 28°C), nighttime temperature (10, 20 or 28°C), daytime light (upper panel) or daytime shade (lower panel) and either control or end-of-day far-red treatment (EOD FR). The left- and right-hand side of the symbols indicate daytime and nighttime temperatures, respectively. Box plots show median, interquartile range 1-3 and the maximum-minimum interval of 4 biological replicates (see Table S1A for detailed statistics). **(B)** Hypocotyl length increment measured in Col-0 seedlings during the night (starting at ZT= 10h) as affected by four combinations of daytime temperature (10 or 28°C) and nighttime temperature (10 or 28°C). All seedlings received shade during the day and EOD FR. Data are means ± SE of nine biological replicates for each time point. The interaction between nighttime and daytime temperatures persists beyond ZT= 14 (see Table S1B for detailed statistics).

To obtain a detailed kinetics of nighttime growth, we exposed the seedlings to two different temperatures during the day (10 and 28°C) and two different night temperatures (10 and 28°C) in all four combinations. Hypocotyl growth rate responded to current and previous temperature conditions (Fig. 1B, Table S1B). After the initial 4 h of the night, hypocotyl growth proceeded at a constant rate, indicating that the transition interval had ended by then. However, growth rate remained significantly affected by previous temperatures beyond that point (Fig. 1B). In effect, according to a multiple regression analysis, hypocotyl length increase between ZT= 14 h and ZT= 22 h depended on time (coefficient ±SE, 4.4 E-03 ±1.4E-03, p=0.0025), the interaction between time and night temperature (0.04±1.4E-03, p <0.0001), and the interaction with daytime temperature (0.01±1.4E-03, p <0.0001). These observations indicate that the daytime cue remains stored in the system, persistently affecting growth. In plants that experienced a cold day, a warm night did not increase hypocotyl growth to the rates exhibited by plants already exposed to warmth during the day. Conversely, in plants that experienced a warm day, a cold night did not decrease hypocotyl growth to the rates exhibited by plants already exposed to cold temperature during the day.

### Nighttime gene expression depends on daytime temperatures

Prompted by these interaction effects we investigated whether nighttime gene expression depended on daytime temperatures. We used transgenic lines bearing the *pPIL:LUC* or *pIAA19:LUC* promoter-reporter fusions in seedlings grown at two different temperatures during the day (10 and 28°C) and two different night temperatures (10 and 28°C) in all four combinations. For both promoters, the activities in seedlings transferred from 10 to 28°C did not reach the levels observed in seedlings exposed to 28°C day and night; and in the seedlings transferred from 28 to 10°C did not drop to the levels observed for those exposed to 10°C day and night (Fig. 2A-B, Table S1C-D). Noteworthy, a cold night did not cause any reduction in the activity of the *pPIL:LUC* promoter after a warm day. Therefore, not only growth but also gene expression showed effects of daytime temperature that persisted during the night.

**Figure 2.**
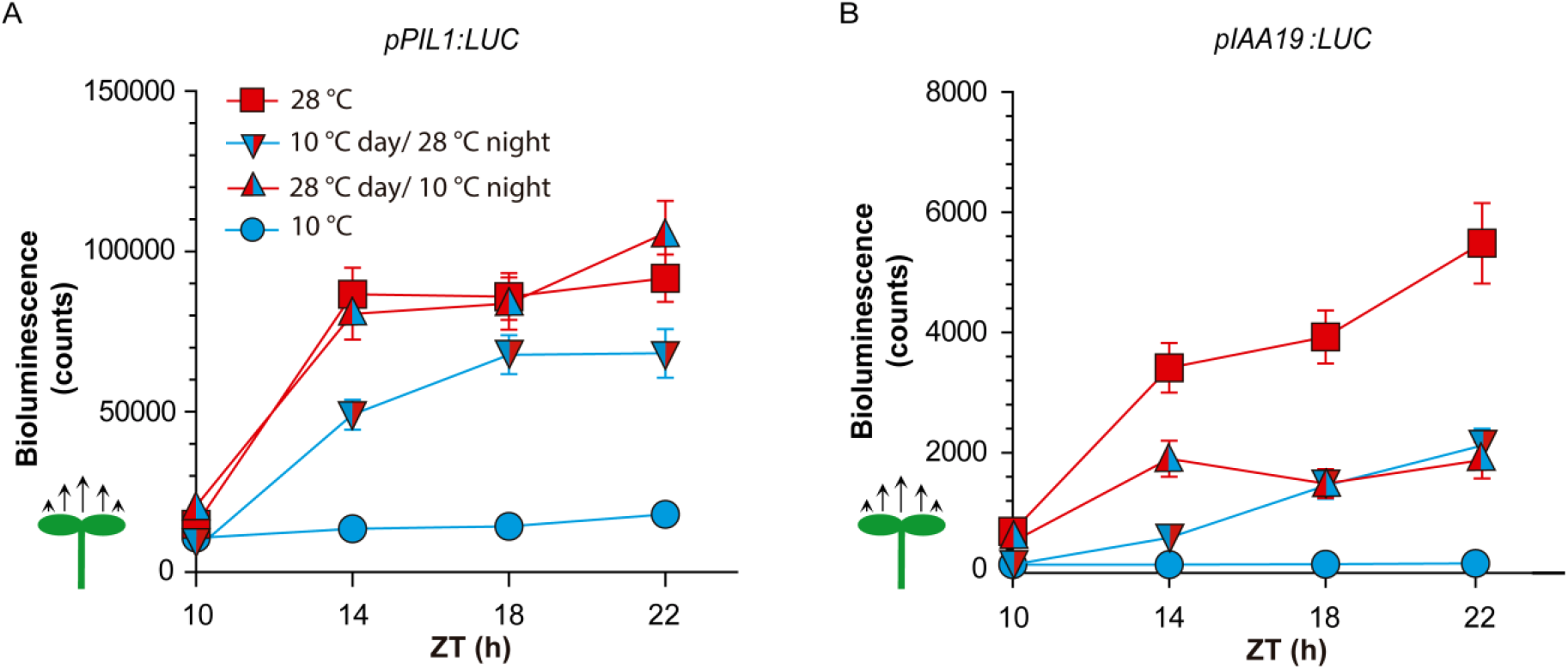
Daytime temperatures affect nighttime gene expression. **(A)** and **(B)** Time course of luciferase activity driven by the *pPIL1:LUC* (A) or *pIAA19:LUC* (B). Luminescence was recorded during the night (starting at ZT= 10h) as affected by four combinations of daytime temperature (10 or 28°C) and nighttime temperature (10 or 28°C). All seedlings received shade during the day and EOD FR. Data are means ± SE of three plates with 96 seedlings (see Tables S1C-D for detailed statistics).

### Genetic requirement of daytime temperature effects on nighttime growth

We compared the WT to different mutants to gain insight into the genetic requirements for the effect of daytime temperature on nighttime growth. We grew seedlings at 28°C or 10°C and exposed them to shade during daytime, followed by EOD FR. In a first set of experiments, we exposed all the seedlings to 28°C during the night. The WT showed faster growth during the night when subjected to the warmer temperature during daytime (Fig 3A-B, Table S1E). The *pif4, pif7* and *hfr1* mutants showed a reduced response to daytime temperature and the *cop1, elf3* and *hy5* mutants showed an inverted response. The *pif5* mutation did not reduce the effect of daytime temperature and actually partially rescued the defect of *pif4* and *pif7* (Fig. S1, Table S1E). In a second set of experiments, we exposed all the seedlings to 10°C during the night. Loss of effects of daytime temperature were observed in the *cop1, pif4, pif5, pif7* and *hy5* mutants (Fig 3C-D, Table S1F).

**Figure 3.**
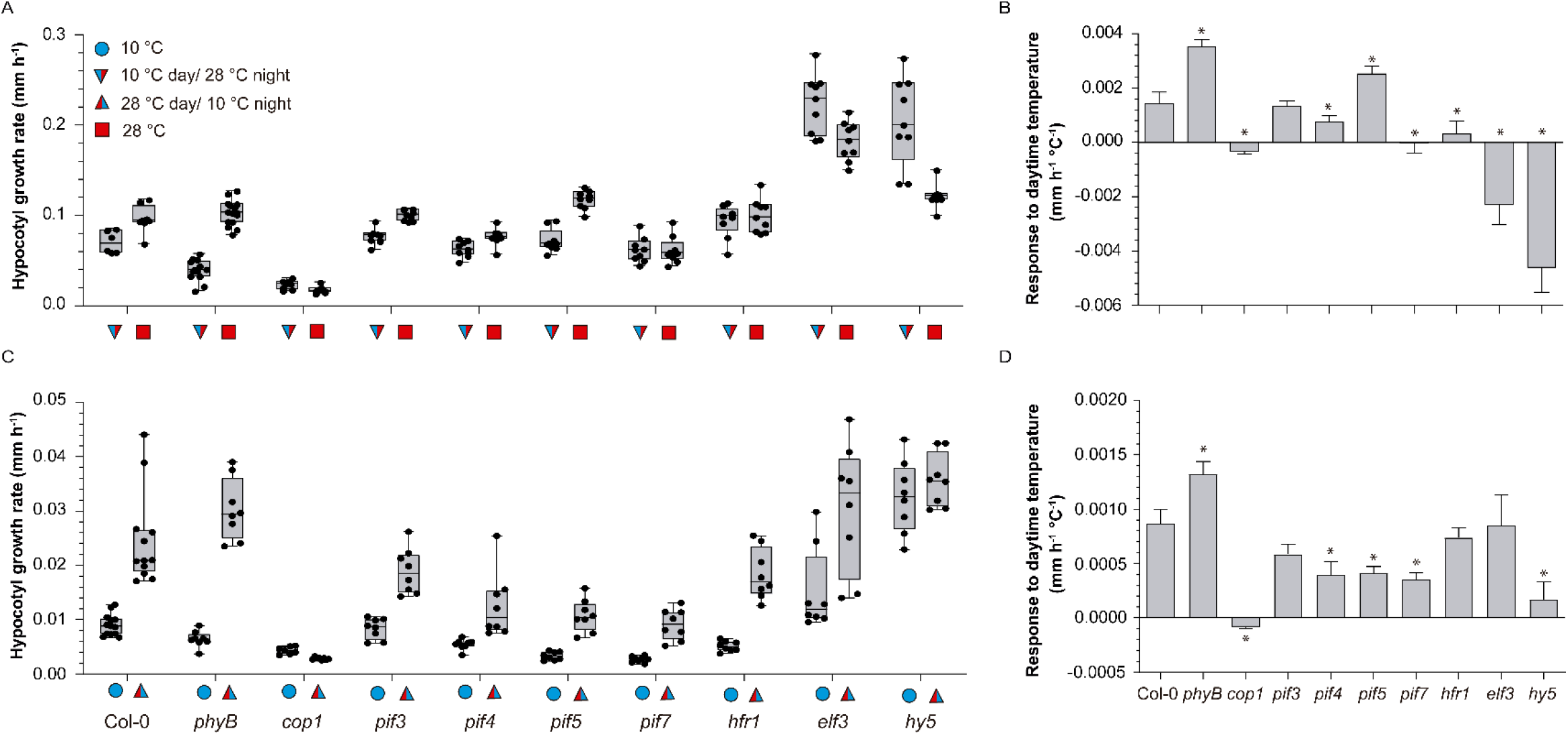
Genetic requirements of the effects of daytime temperature on nighttime growth. **(A)** and **(C)** Hypocotyl growth rate measured in seedlings of the indicated genotypes during the night (ZT 10h to ZT 24h) as affected by two different daytime temperatures (10 or 28°C). **(B)** and **(D)** Slope of the responses to daytime temperature (linear regression analysis of the data in A and C, respectively). All seedlings received shade during the day and EOD FR. Night temperature was 28°C (A-B) or 10°C (C-D). Box plots show median, interquartile range 1-3 and the maximum-minimum interval of 8-12 biological replicates (see Table S1 E-F for detailed statistics). In B and D, the asterisks indicate significant differences with Col-0 according to *t*-tests (*, p<0.05).

### Memory of daytime temperature in the status of signalling components

The *pif4, pif7, hfr1, cop1, elf3* and *hy5* mutants did not show the normal enhancement of nighttime growth by warm daytime temperatures observed in the WT. Therefore, we investigated the status of these signalling components by 4 h into the night (ZT= 14 h, 28°C) in seedlings exposed to contrasting daytime temperatures (10 and 28°C).

Warm temperatures increase the nuclear levels of PIF4 (Legris et al., 2017) and COP1 (Park et al., 2017) and decrease the nuclear levels of HY5 (Romero-Montepaone et al., 2021). We used confocal microscopy to analyse the nuclear fluorescence of *pPIF4:PIF4-GFP, pHY5:HY5-YFP* and *p35S:YFP-COP1* transgenic lines. At 28°C, nuclear abundance of PIF4 and COP1 in the night was increased and that of HY5 was reduced in the hypocotyl cells of seedlings that had received 28°C compared to 10°C during the day (Fig. 4A-F).

**Figure 4.**
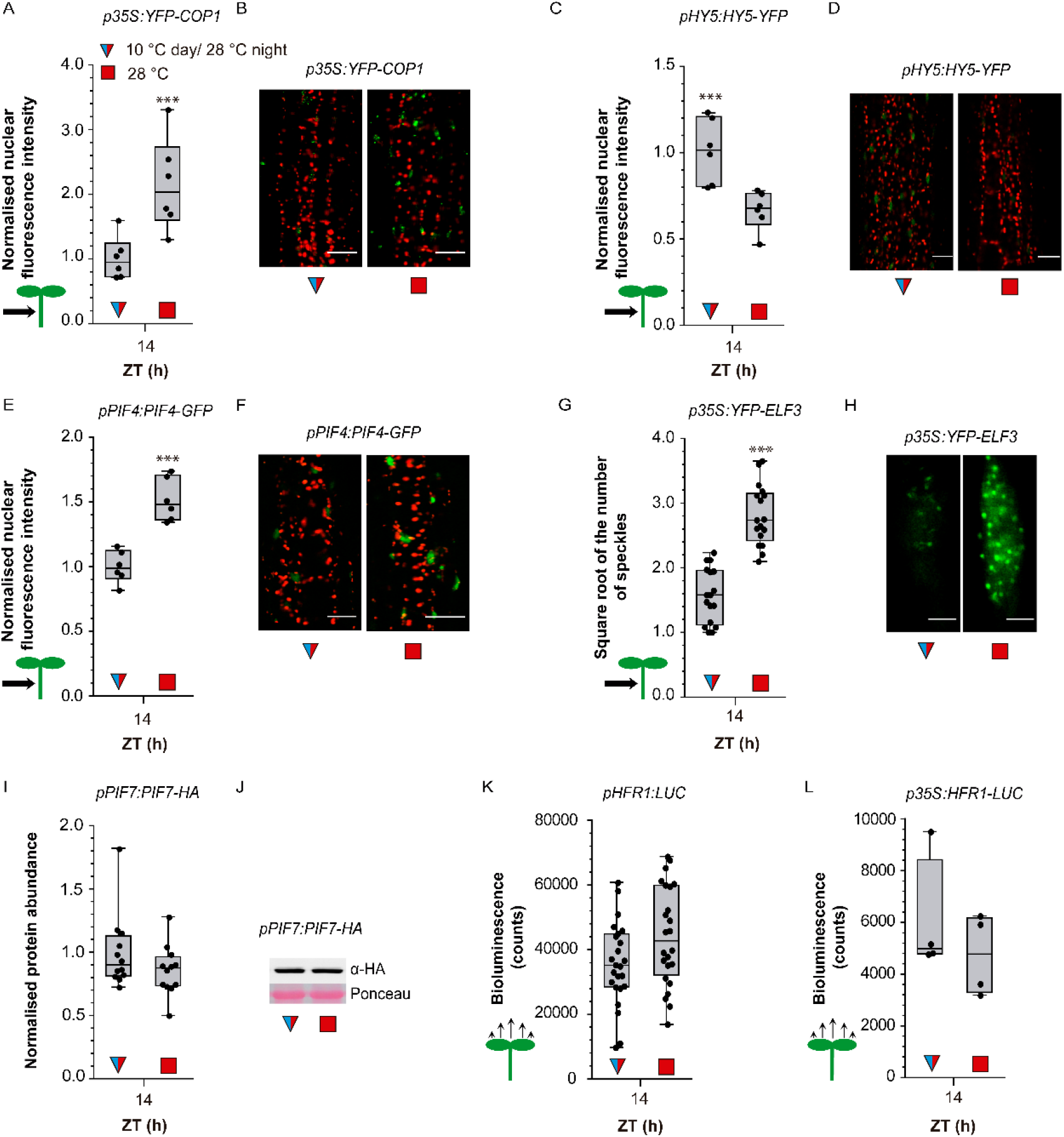
Daytime temperatures affect nighttime signalling status. **(A-B)** Nuclear fluorescence driven in hypocotyl cells by *p35S:YFP-COP1*. **(C-D)** Nuclear fluorescence driven in hypocotyl cells by *pHY5:HY5-YFP*. **(E-F)** Nuclear fluorescence driven in hypocotyl cells by *pPIF4:PIF4-GFP*. **(G-H)** Number of ELF3 nuclear speckles (square root-transformed data) in the hypocotyl cells of the line expressing the *p35S:YFP-ELF3* transgene. **(I-J)** Abundance of PIF7 in seedlings bearing *pPIF7:PIF7-HA*. **(K-L)** Luciferase activity driven by *pHFR1:LUC* (K) or *p35S:HFR1-LUC* (L). The seedlings received shade and either 10°C or 28°C during the day, EOD FR and 28°C during the night. Box plots show median, interquartile range 1-3 and the maximum-minimum interval of 6 (A, C and E), 16 (G), 12 (I), 25 (K), 4 (L) biological replicates. Representative confocal (B, D, F, H) or protein blot (J) images. Scale bar = 35 µm (B, D, F) and 2.61 µm (H). Asterisks indicate significant differences in *t-*tests (*, p<0.05; **, p<0.01; ***, p< 0.001).

Warm temperatures were described to increase (Jung et al., 2020) or decrease (Ronald et al., 2021) the formation of nuclear speckles containing ELF3. Since overexpression of ELF3 has no effect on temperature responsivness compared to the WT (Thines and Harmon, 2010; Jung et al., 2020), we used the *p35S:YFP-ELF3* line to facilitate the quantitative analysis of the ELF3 speckles (the same line used by Ronald et al., 2021). We observed a significantly higher number of ELF3 speckles in the seedlings exposed to 28°C than in seedlings exposed to 10°C during the day (Fig. 4G-H).

Warm temperatures increase PIF7 protein levels (Fiorucci et al., 2020; Chung et al., 2020) and enhance HFR1 protein stability (Romero-Montepaone et al., 2021). However, protein blot analyses of seedlings sampled after four hours into the 28 °C night did not reveal differences in PIF7 protein abundance caused by contrasting daytime temperatures (Fig. 4I-J). Similarly, in luminometer readings, we did not observe significant effects of daytime temperature on nighttime *HFR1* promoter activity or HFR1 stability (Fig. 4K-L).

PIF4 and COP1 promote hypocotyl growth whereas HY5 inhibits hypocotyl growth during the night (e.g. Fig. 3). Cold days decreased PIF4 and COP1 nighttime activities and increased HY5 nighttime activities (Fig. 4A-F), suggesting that PIF4, HY5, and COP1 convey daytime temperature information to nighttime growth. This would also be the case for ELF3 if enhanced speckle formation correlates with reduced activity, a link analysed in further detail below. Conversely, the short-term memory of daytime temperature requires PIF7 and HFR1 (Hornitschek et al., 2009; Sandi Paulišić et al., 2021), but apparently, these factors do not carry prior temperature information because their levels during the night showed no influence of daytime conditions (Fig.4I-L).

### Nighttime PIF4 and HY5 kinetics account for growth responses

Since ELF3 and COP1 control hypocotyl growth partially via PIF4 and HY5 (Gangappa and Kumar, 2017; Park et al., 2017; Nusinow et al., 2011; Box et al., 2015), we analysed in further detail the nighttime kinetics of nuclear abundance of these two transcription factors by using confocal microscopy of hypocotyl cells. The seedlings exposed to 28°C during the day, compared to 10°C, during the day, initiated the night with more PIF4 and less HY5 in the nucleus of their hypocotyl cells and these differences persisted at least during several hours (Fig. 5A-D). The seedlings that were transferred at the beginning of the night from 10°C to 28°C reached similar PIF4 or HY5 nuclear levels as those kept at 28°C during daytime by 8 h after the temperature shift (ZT= 18 h, Fig. 5A-D). In seedlings transferred from 28°C to 10°C, similar HY5 nuclear levels as in the seedlings already exposed to 10°C during daytime were only observed 12 h later (ZT= 22 h, Fig. 5C-D). Noteworthy, 28°C to 10°C shift at the beginning of the night caused almost no PIF4 response (Fig. 5A-B). Taken together, these results indicate that the nighttime kinetics of PIF4 and HY5 depend on both nighttime and daytime temperature (Table S1G-H). Differences in their protein levels caused by the contrasting temperatures during the day extended several hours into the night, either due to a slower transitions between the levels typical of day and night temperatures (PIF4 from 10°C to 28°C and HY5 in both directions) or a nearly complete insensitive response (PIF4 from 28°C to 10°C).

**Figure 5.**
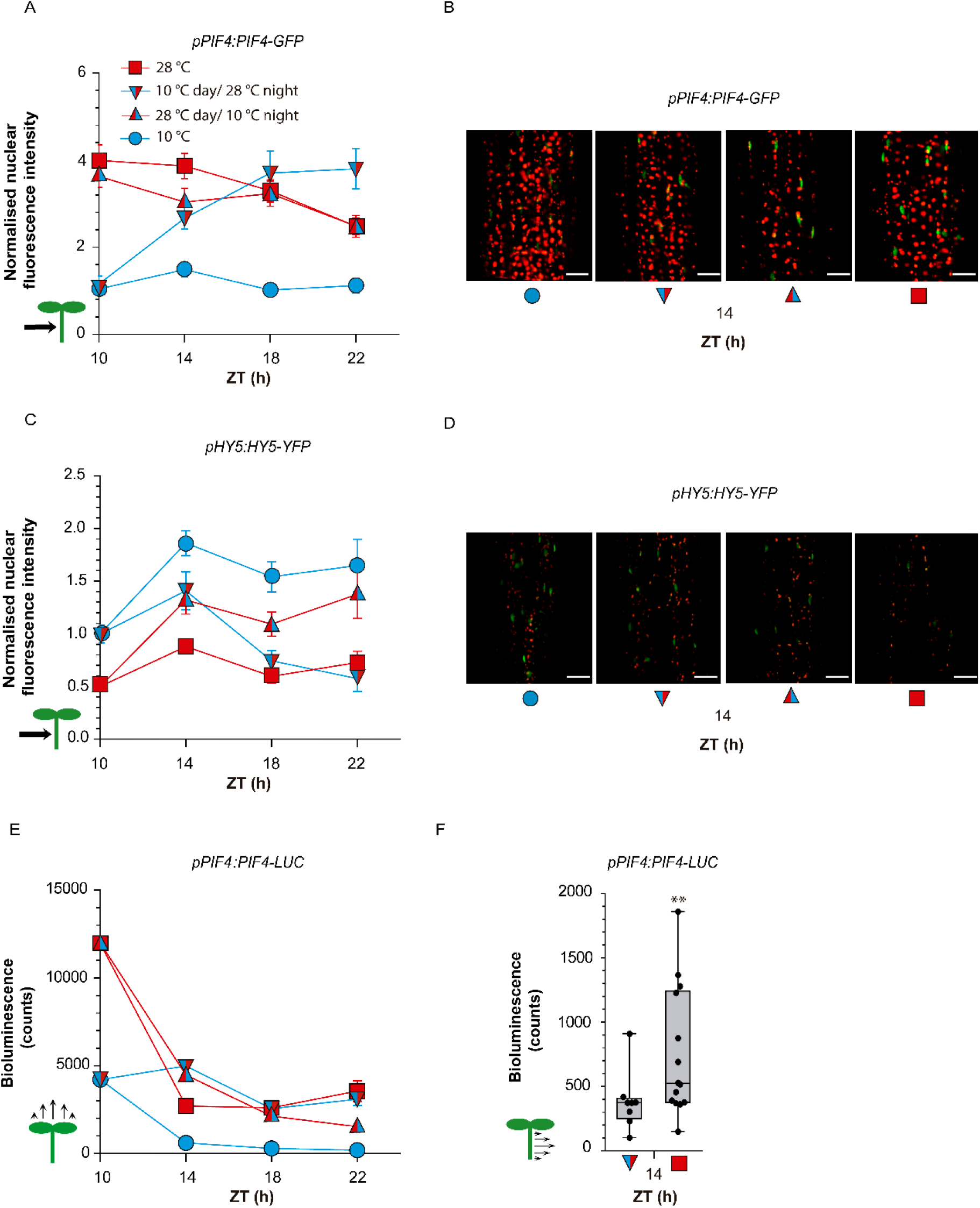
Daytime temperature affects nuclear levels of PIF4 and HY5 during the night. **(A-D)** Nighttime kinetics of nuclear fluorescence intensity driven in hypocotyl cells by *pPIF4:PIF4-GFP* (A-B) or *pHY5:HY5-YFP* (C-D). **(E)** Nighttime kinetics of luminescence driven by *pPIF4:PIF4-LUC* in entire seedlings. **(F)** Luminescence driven by *pPIF4:PIF4-LUC* in isolated hypocotyls harvested at ZT= 14 h. Confocal images or luminescence values were recorded during the night (starting at ZT= 10h) as affected by four combinations of daytime temperature and nighttime temperature. All seedling received shade during the day and EOD FR. (A, C, E) Data are means ± SE of three biological replicates. (F) Box plots show median, interquartile range 1-3 and the maximum-minimum interval of 10-14 biological replicates. (B, D) Representative confocal images at ZT= 14h. Scale bar = 35 µm. See Table S1G-I for detailed statistics of (A, C, E). In (F), the asterisks indicates significant differences in *t-*test (**, p<0.01).

Genetic studies indicated that effects of daytime temperature effects depend on PIF4 and HY5 (Fig. 3). Therefore, we explored the quantitative association between the growth rate kinetics of growth rate and that of PIF4 and HY5 nuclear levels by using multiple regression analysis across three temporal phases (ZT 10-14 h, 14-18 h and 18-22 h). For each temporal phase, we averaged the PIF4- or HY5-fluorescence values obtained by confocal microscopy at the beginning and the end of the phase (Fig. 5A and C) as explanatory variables. The model accounted for a significant proportion of the variability in growth rate (adjusted R^2^=0.80, P <0.0001). Both PIF4 and HY5 levels contributed significantly (coefficient ±SE, PIF4: 2.1E-03 ±1.5E-04, p <0.0001; HY5: -1.0E-03 ±3.2E-04, p= 0.016), indicating that the information that provided by PIF4 and HY5 dynamics is important and not redundant.

We also investigated the abundance of the PIF4 protein during the night by using transgenic lines bearing *pPIF4:PIF4-LUC*. As in the case of confocal microscopy analysis, there was little difference in PIF4 protein levels if the plants continued at 28°C or were shifted from 28°C to 10°C at the beginning of the night (Fig. 5E, Table S1I). Also resembling the hypocotyl pattern, PIF4 remained low throughout the night in plants grown at 10°C during the day and the night, and increased when plants were transferred from 10 °C to 28°C at the beginning of the night (Fig. 5E), although this response was faster than in the confocal studies. The upwards cotyledon signal dominates luminescence readings of entire seedlings; therefore, we used the line bearing the *pPIF4:PIF4-GFP* transgene to evaluate nuclear levels of PIF4 in cotyledon cells. The results confirmed elevated levels of PIF4 during the day (see also Stavang *et al*., 2009) and showed that these differences had already been inverted by 4 h into the night (Fig. S2). The bioluminescence analysis of PIF4 levels in isolated hypocotyls of the *pPIF4:PIF4-LUC* line actually confirmed that the shift to 28°C at the beginning of the night was not enough to achieve the levels detected in this organ in seedlings grown at 28°C during the day (Fig. 5F).

Given the faster change in PIF4 levels in the cotyledons, we investigated whether the growth of this organ responds to daytime temperature in a PIF4-dependent manner. The area of the cotyledons increased more during the night at 28°C if the seedlings were exposed during daytime to 10°C (mean ±SE, n= 40, 4.0 E-03 ± 4.0 E-04 mm^2^), than when daytime temperature was 28°C (2.7 E-03 ± 2.1 E-04 mm^2^, P =0.005). This memory effect was absent in the *pif4* mutant. Compared to the WT, *pif4* showed an enhanced cotyledon expansion with 28°C daytime temperature (4.3 E-03 ± 4.8 E-04 mm^2^, P =0.0017), similar to that observed at 10°C (4.2 E-03± 4.8 E-04 mm^2^) (Huq and Quail, 2002). Therefore, differences in PIF4 levels in the cotyledons generated by 28°C compared to 10°C during the day, in spite of showing a shorter persistence during warm nights, convey daytime temperature information to nighttime growth of this organ.

### *PIF4* promoter activity accounts for PIF4 dynamics during the night

Nighttime kinetics of *PIF4* promoter activity conserved the key features observed for the PIF4 protein kinetics. It depended on nighttime and daytime activity and showed asymmetric responses to temperature shifts (Fig. 6A, Table S1L). The stronger *PIF4* promoter activity established by 28°C compared to 10°C during the day was largely irreversible during the night, as transfer to 10°C did not reduce this activity below the levels of 28°C controls (Fig. 6A). Conversely, the seedlings transferred from 10°C to 28°C at the beginning of the night did increase the activity of the *PIF4* promoter (Fig. 6A). Luminescence readings driven by promoter and protein fusions showed strong correlation (Fig. 6B), supporting a major role of *PIF4* promoter activity in the control of nighttime PIF4 levels.

**Figure 6.**
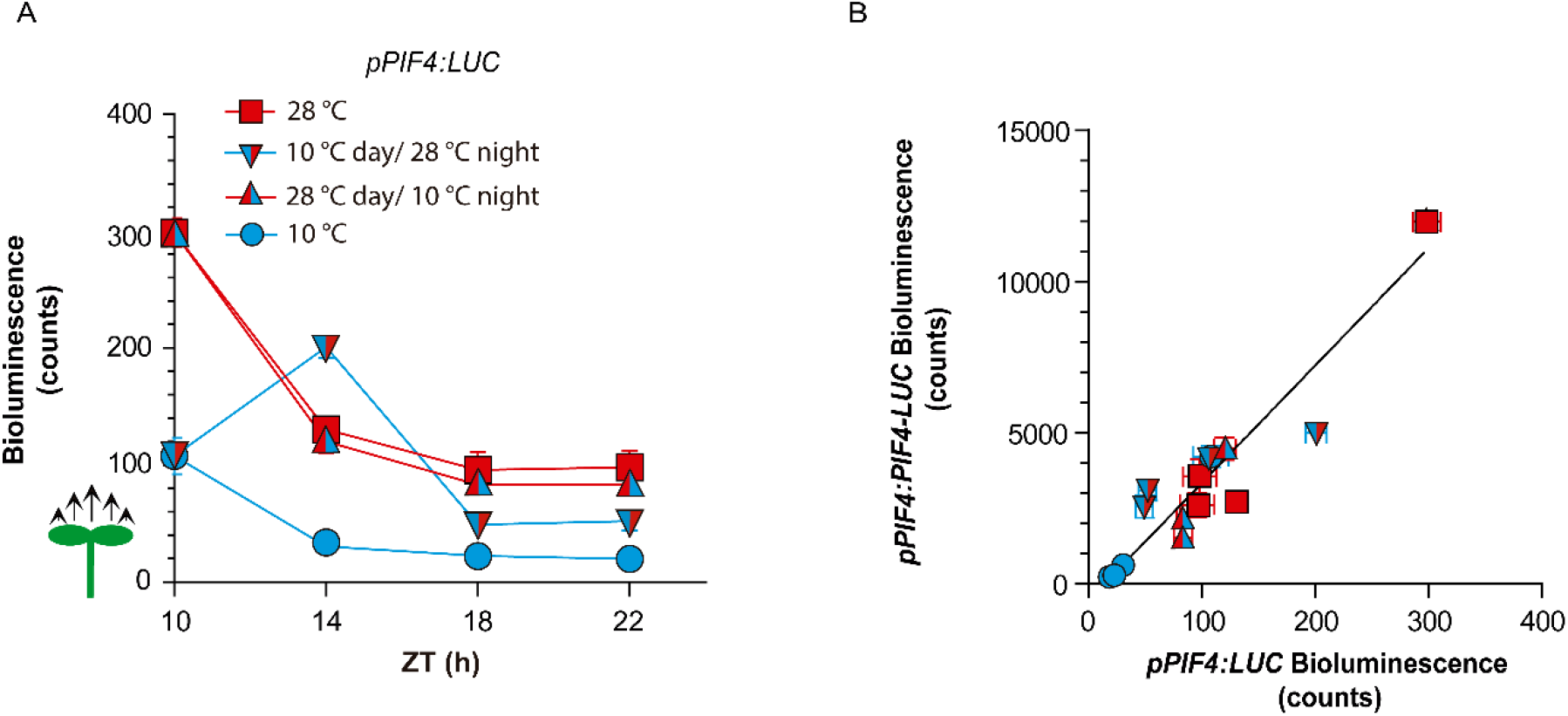
Changes in PIF4 promoter activity account for nighttime PIF4 protein dynamics. **(A)** Nighttime kinetics of luminescence driven by *pPIF4:LUC* in entire seedlings. Luminescence was recorded during the night (starting at ZT= 10h) as affected by four combinations of daytime temperature (10 or 28°C) and nighttime temperature (10 or 28°C). All seedling received shade during the day and EOD FR. Data are means ± SE of three plates with seedlings. See Table S1J-K for detailed statistics. **(B)** Correlation between luminescence driven by *pPIF4:PIF4-LUC* (from 5E) and *pPIF4:LUC* (from A) in the Col-0 background. The correlation is significant at P<0.0001.

The nighttime patterns of PIF4 protein and *PIF4* promoter activities showed two features. The first feature is that daytime differences persisted at least several hours into the night. The second feature is the asymmetric response to temperature shifts in contrasting directions (increase compared to decrease in temperature). This asymmetric response to a variable when it either increases or decreases its values is typical of hysteretic systems (Davies, 2017; Jiang and Hao, 2021).

### Daytime generation of night differences in PIF4 requires ELF3

The differences in PIF4 nuclear levels (Fig. 5A, C) and *PIF4* promoter activity (Fig. 6A) that persisted during the night were already present in the cotyledons and hypocotyls at the end of the day. We therefore investigated the processes involved in their generation during the photoperiod. Luminescence readings revealed that the abundance of PIF4 protein (Fig. 7A) and the activity of the *PIF4* promoter (Fig. 7B) abruptly increased early in the morning, when the seedlings were first transferred from 20°C to 28°C, compared to 10°C. Initial changes induced by such contrasting temperatures slightly narrowed down during the course of the day but were still present at the beginning of the night. The early increase in PIF4 protein signal at warm temperatures was more intense than that in *PIF4* promoter activity (cf. Fig 7A and B at 4 h), which is reflected by a higher protein / promoter activity ratio (Fig. 7C). The protein blot results using a line bearing the *p35S:PIF4-HA* transgene are consistent with a higher stability of PIF4 at 28°C than at 10°C early in the morning (Fig. S3A-B). However, later on in the day, PIF4 protein levels increased in the plants exposed to 10°C without a concomitant increment in *PIF4* promoter activity, causing a higher protein / promoter activity ratio in plants exposed to cooler temperatures (Fig. 7C). Therefore, the differences in PIF4 protein levels at the beginning of the night were largely due to temperature effects on *PIF4* promoter activity, with only transient post-transcriptional effects.

**Figure 7.**
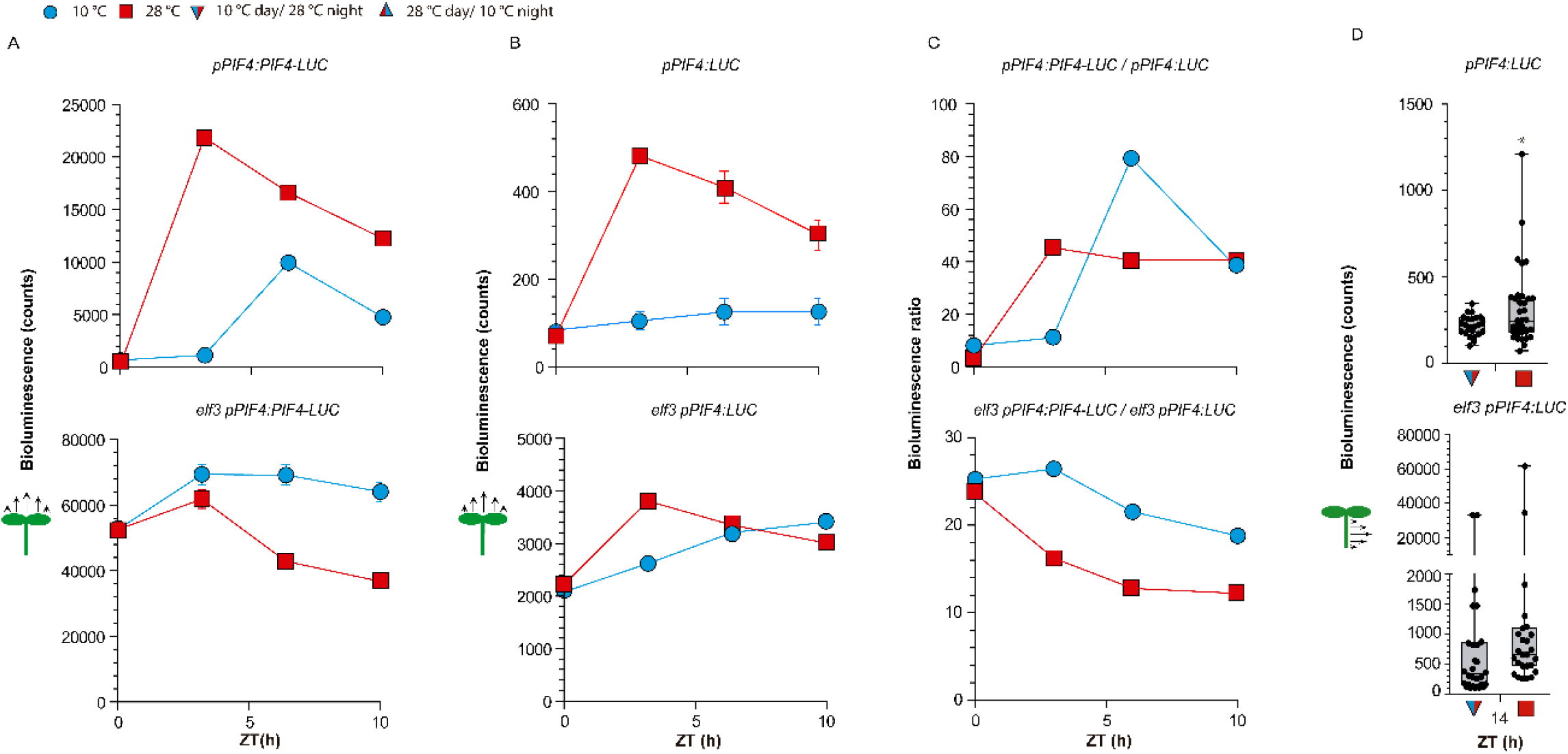
Temperature effects on end-of-day *PIF4* promoter activity and PIF4 protein abundance require ELF3. **(A-B)** Time course of daytime luciferase activity driven by *pPIF4:PIF4-LUC* (A) or *pPIF4:LUC* (B) in the Col-0 and *elf3-8* background, as affected by 10°C or 28°C. **(C)** Ratio between the luciferase signals driven by *pPIF4:PIF4-LUC* and *pPIF4:LUC*. **(D)** Luminescence driven by *pPIF4:LUC* in isolated hypocotyls harvested from Col-0 and *elf3-8* seedlings at ZT= 14 h. All seedling received shade during the day and EOD FR. (A-B) Data are means ± SE of 3 plates with seedlings. (D) Box plots show median, interquartile range 1-3 and the maximum-minimum interval of 24-30 biological replicates, asterisk indicates significant differences in *t-*tests (*, p<0.05). See Table S1L-M for detailed statistics of (A-B).

We investigated whether differences in PIF4 observed at the end of the day require ELF3. Experiments using the lines bearing the *pPIF4:LUC* transgene in the *elf3* background indicated that, early in the photoperiod, warm temperature promoted *PIF4* activity even in the absence of ELF3 (Fig 7B), implying the action of other transcriptional regulators. However, temperature-induced changes in *PIF4* promoter activity showed an absolute requirement of ELF3 during the rest of the photoperiod (Fig 7B). During warm afternoons *PIF4* promoter activity declined, even in the *elf3* background (Figs 7B), and this might reflect an effect of GIGANTEA (Anwer et al., 2020).

Since luminometer readings of entire seedlings reflect *PIF4* promoter activity mainly in the cotyledons, we conducted experiments with isolated hypocotyls; i.e., the organ where differences in PIF4 are more persistent during warm nights (*cf* Figs 5C and E). The memory of daytime temperatures of the *PIF4* promoter was also absent in the hypocotyl in the *elf3* background (Fig. 7D).

### Kinetics of ELF3

We investigated the dynamics of ELF3 under the conditions where it controlled *PIF4* promoter activity. Under warm temperature, the formation of speckles decreased during the morning and increased strongly during the afternoon (Fig. 8A and C). In addition to *PIF4*, ELF3 negatively regulates *TOC1* and positively regulates the *LHY* and *CCA1* promoters (Thines and Harmon, 2010; Fehér et al., 2011; Herrero et al., 2012; Ezer et al., 2017), in all cases through direct binding of these promoters. The activity of the *TOC1* promoter increased whilst that of the *LHY* and *CCA1* promoters decreased in response to warm temperature (Fig. S4). In particular, *TOC1* did not show responses during the morning. Taken together with the pattern of *PIF4*, these results indicate a negative correlation between ELF3 speckle formation and ELF3 activity.

**Figure 8.**
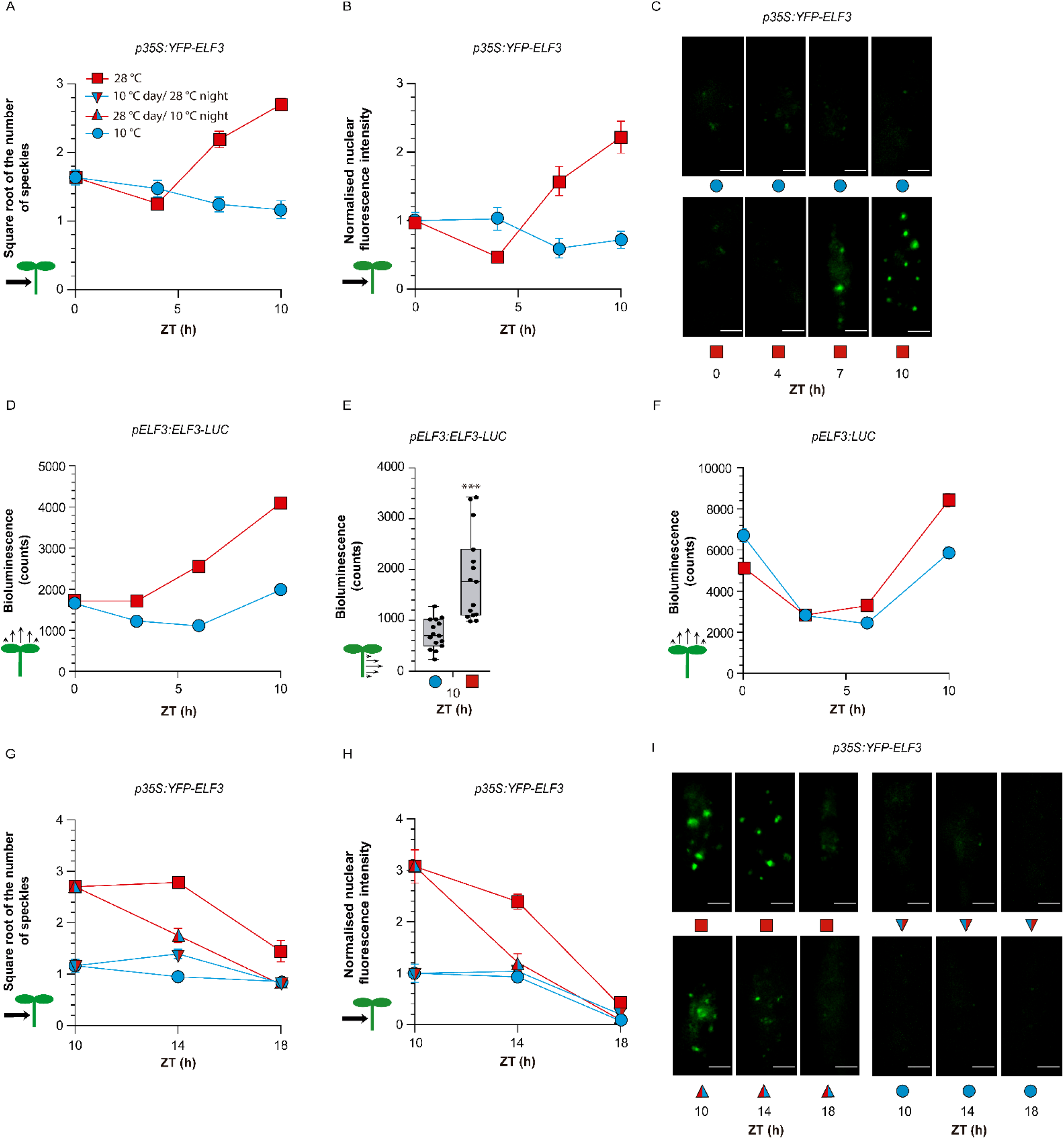
ELF3 dynamics under contrasting temperatures. **(A-C)** Number of ELF3 nuclear speckles (A), nuclear fluorescence intensity (B) and representative confocal images (C) from hypocotyl cells of seedlings expressing *p35S:YFP-ELF3*. **(D)** Luciferase activity driven by *pELF3:ELF3-LUC* in entire seedlings. **(E)** Luciferase activity driven by *pELF3:ELF3-LUC* in isolated hypocotyls. **(F)** Luciferase activity driven by *pELF3:LUC* in entire seedlings. **(G-I)** Number of ELF3 nuclear speckles (square root-transformed data, G), nuclear fluorescence intensity (H) and representative confocal images (I) from hypocotyl cells of seedlings expressing *p35S:YFP-ELF3*. All seedling received shade during the day and EOD FR. (A-B and G-H) Data are means ± SE of 20 biological replicates. (C and I) Representative confocal images. Scale bar = 2.61 µm. (D and F) Data are means ± SE of three plates with seedlings. (E) Box plots show median, interquartile range 1-3 and the maximum-minimum interval of 15 biological replicates. See Table S1N-S for detailed statistics of (A-B, D, F-H). In E, asterisks indicate significant differences in *t-*test (***, p< 0.001).

During the morning, warm temperature decreased ELF3 nuclear abundance (Fig. 8B) and the bioluminescence signal driven by the *pELF3:ELF3-LUC* transgene (Fig. 8D). During the afternoon, warm temperature increased both features (Fig. 8B, D). Harvested hypocotyls also showed elevated nuclear levels of ELF3 at 28°C (Fig. 8E). Conversely, bioluminescence driven by the *pELF3:LUC* transgene increased only slightly at 28°C (Fig. 8F). These results suggest a post-transcriptional control of the ELF3 protein levels, which might involve increased stability within the speckles.

As described above (Fig. 4G-H), during the night, the shift from 10 °C to 28 °C did not increase the number of ELF3 speckles to the levels observed in seedlings already exposed to 28 °C during the day. Actually, the number of speckles decreased during the night in all temperature conditions (Fig. 8G and I). However, such a decrease does not reflect enhanced ELF3 activity because a steady drop in ELF3 protein levels accompanied the decrease in speckles (Fig. 8H and I).

### Hysteresis in *PIF4* promoter activity

In all the experiments described above, temperature shifts coincided with end of the day. However, under natural conditions temperature fluctuations can occur during the photoperiod. We therefore investigated whether the response pattern of the *PIF4* promoter conserved in the day the behaviour observed during the night. For this purpose, we exposed the seedlings to 10°C or 28°C and transferred them at ZT= 4h, either from 10°C to warmer temperatures (15, 20, 25 or 28°C) or from 28°C to cooler temperatures (25, 20, 15 or 10°C), whilst the controls remained at 10°C or 28°C. We harvested the seedlings 3 hours after the temperature shift, i.e., still within the photoperiod (ZT= 7h). Fig. 9A shows a memory of the previous temperature because for most afternoon temperatures (abscissas) warmer morning temperatures yielded higher *PIF4* promoter activities for most afternoon temperatures (abscissas). This memory is entirely associated to hysteresis in promoter activity, demonstrated by a shift in sensitivity in the way up as compared to the way down.

**Figure 9.**
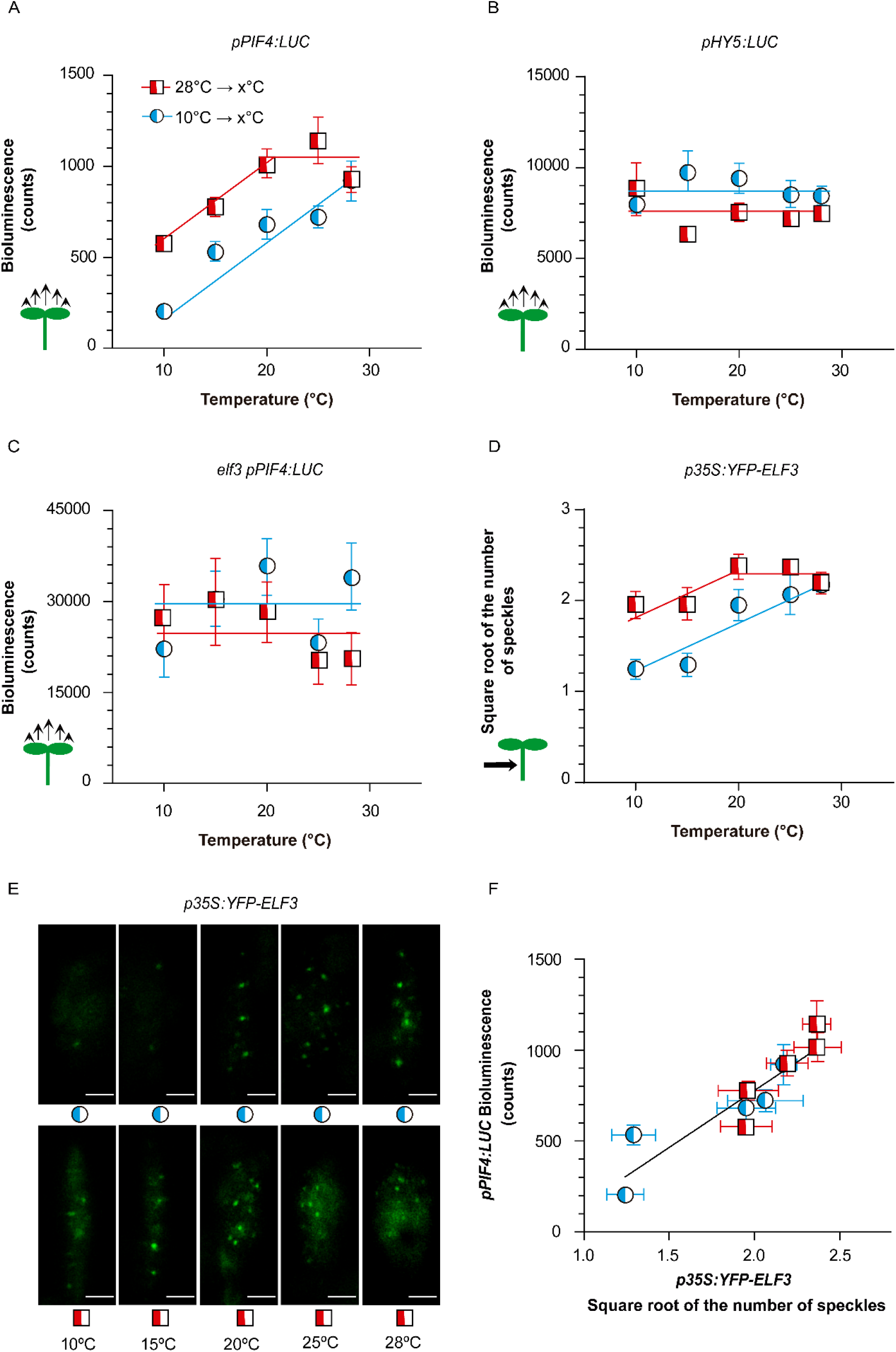
Hysteresis of ELF3 drives hysteresis in *PIF4* promoter activity. **(A-C)** Response of the luminescence driven by *pPIF4:LUC* (A and C) or *pHY5:LUC* (B) in the Col-0 (A-B) or *elf3* (C) backgrounds to increasing or decreasing temperature. **(D-E)** Response of the number of ELF3 nuclear speckles (square root-transformed data) from hypocotyl cells of seedlings expressing *p35S:YFP-ELF3* to increasing or decreasing temperature. **(F)** Correlation between luminescence driven by *pPIF4:LUC* (from A) and the number of speckles driven by *p35S:YFP-ELF3* (from D). The seedlings were exposed to white light at either 10°C or 28°C and 4 h after the beginning of the photoperiod were transferred to the temperature indicated in abscissas (including controls that remained at 10°C or 28°C). Luminescence (A-C) or confocal images (D and E) were taken 3 h later. Data are means ± SE of 3 plates with seedlings (A-C) or 15 biological replicates (D and E). (E) Representative confocal images. Scale bar = 2.61 µm. In F, the correlation is significant at P =0.0002. See Table S1T-X for detailed statistics.

We simultaneously analysed the activity of the *HY5* promoter, which remained largely unresponsive to the temperature shifts in any of the two directions (Fig. 9B). This indicates that different mechanisms mediate the persistent effects of previous temperature on PIF4 and HY5 nuclear levels. Compared to the WT, the analysis of seedlings bearing *pPIF4-LUC* in the *elf3* mutant background showed a completely distorted pattern of response to temperature (Fig. 9C). This indicates that the hysteresis pattern depends on ELF3. We therefore investigated the response of ELF3 itself.

### Hysteresis in ELF3 speckle formation

We then analysed the formation of the ELF3 speckles in response to increasing or decreasing temperatures during the day, in the same conditions used to investigate the hysteresis of the *PIF4* promoter. The seedlings exposed to 28°C showed more speckles than those exposed to 10°C (Fig. 9D). Noteworthy, the response curve showed a strong shift in sensitivity, which was higher in the way up than in the way down. Therefore, ELF3 sub-nuclear location showed in itself the hysteretic pattern. The activity of the *PIF4* promoter strongly correlated with the number of ELF3 speckles formed under the same conditions (Fig. 9E), supporting a link between speckle formation and reduced ELF3 activity.

Compared to 10°C, 28°C did not increase the number of speckles during the first 4 h of the day (Fig. S5, see also Fig. 8A). Therefore, all the differences observed in Figure 9D originated during the afternoon (i.e., between 4 and 7h; Fig. S5). This means that the speckles appear rapidly during the afternoon, but not in the morning (Fig. S5), suggesting that other components required for their formation of the speckles become available in the afternoon. Furthermore, the additional speckles formed between 4 and 7 h, even in seedlings exposed to during the first 4 h at 28°C and then transferred to 10°C. This demonstrates that warmth induction of ELF3 speckle formation persisted in the cold until the additional putative components of the speckles became available. The persistence of a warmth-induced state of ELF3 is consistent with the occurrence of hysteresis.

## Discussion

The results reported here demonstrate that the control of hypocotyl growth and gene expression during the night respond not only to the current temperature environment but also to the temperature experienced during the preceding photoperiod (Figs 1-2). Therefore, there is a nighttime memory of daytime temperature.

PIF4 and HY5 store daytime temperature information to the control of hypocotyl growth during the night. First, the genetic analysis of the growth response indicated that the nighttime memory of daytime temperatures requires HY5 and PIF4 (Fig.3). Second, both HY5 and PIF4, showed daytime-induced differences in nuclear abundance, which persisted at least 8 h into the night (Figs 4-5) and correlated with the growth rate during the same period. Third, the growth memory also required COP1 and ELF3, two signalling components that stored daytime temperature information (Fig. 4) and control the abundance of HY5 and/or PIF4 (Gangappa and Kumar, 2017; Park et al., 2017; Nusinow et al., 2011; Box et al., 2015). In the case of PIF4, changes in *PIF4* promoter activity largely accounted for the dynamics of nuclear protein levels, both during daytime (Fig. 7A-B) and at night (Fig. 6A-B). Apparent post-transcriptional effects were only transient and changed properly their direction during the course of the photoperiod (Fig. 6C).

One of the features of the nighttime memory of daytime temperatures is the slow transition in the status of PIF4 and HY5. In the plants shifted from 10°C to 28°C at the beginning of the night, the nuclear levels of PIF4 and HY5 took 8 h to achieve the levels observed in the seedlings that were already at the nighttime temperature during the day (Fig. 5A-B). Similarly, in the plants shifted from 28°C to 10°C at the beginning of the night, the nuclear levels of HY5 took 12 h to reach the levels observed in the seedlings that were already at this nighttime temperature during the day (Fig. 5B). The slow kinetics of these responses is intriguing because at least in the case of PIF4 it represents a night-specific feature. In fact, during the day plants transferred from 10°C to 28°C rapidly (< 3 h) reached the high *PIF4* promoter activity observed in the seedlings already exposed hours earlier to 28°C (Fig. S5). Furthermore, contrary to daytime effects into the night, the differences in hypocotyl nuclear levels of PIF4 generated by nighttime temperatures of 10°C compared to 28°C rapidly (<4 h) reverted during the photoperiod (Murcia et al., 2020). This slower nighttime response of PIF4 might relate to the gating activity of TOC1 (Zhu et al., 2016).

A second feature of the nighttime memory of daytime temperatures is that in plants transferred from warm daytime temperatures to cold nights, PIF4 persisted at the high levels observed in the plants that remained in the warmth (Fig. 5A). Therefore, there is a clear asymmetry in the response of PIF4 to a temperature increase or decrease, with a slow rise of PIF4 in the first case and nearly no response in the latter. The PIF4 target promoter *PIL1* also showed asymmetric responses, as a change from low to high temperature caused a significant increase in activity but the opposite modification resulted in a barely any detectable changes (Fig. 2A). The asymmetric sensitivity of *PIF4* promoter activity was not specific to changes in temperature during the night. In fact, the response curve to temperature changes during the afternoon also indicated greater sensitivity in the way up than in the way down (Fig. 9A). This pattern revealed strong hysteresis of *PIF4* promoter activity; i.e., the response to temperature follows a pattern in the forward direction but a different one in the return direction (Davies, 2017; Jiang and Hao, 2021).

ELF3 played a fundamental role in the dynamics of *PIF4* promoter activity. Although during the morning the *PIF4* promoter responded to temperature even in the absence of ELF3, these differences persisted to the beginning of the night and beyond only in the presence of ELF3 (Fig. 7B, D). The hysteretic pattern of the *PIF4* promoter activity also required ELF3 (Fig. 9C).

Warm temperatures reduce evening complex transcriptional activity (Box et al., 2015; Ezer et al., 2017; Silva et al., 2020) but there is some controversy regarding the response to temperature of nuclear ELF3 speckle formation and the function of these sub-nuclear structures. According to Jung *et al*., (2020) warm temperatures induce the formation of speckles containing ELF3 in root cells but according to Ronald *et al*. (2021), warm temperatures reduce the formation of speckles in hypocotyl and root cells. Here we show that both patterns are not mutually exclusive because warm temperature reduced speckle formation in hypocotyl cells during the morning and increased speckle formation during the afternoon (Fig. 8A, S5). The cellular context appears therefore to affect ELF3 speckle formation. This time of day effect would not be mediated by ELF4 (Ronald et al., 2021). There is a negative association between ELF3 activity and *PIF4* expression (Nusinow et al., 2011; Box et al., 2015; Raschke et al., 2015; Press et al., 2016). Under our conditions, ELF3 speckle formation correlated with enhanced *PIF4* promoter activity (Fig. 9F) and hence reduced ELF3 activity. Membrane-less compartments were linked to changes in the stability of their components (Kim et al., 2021; Emenecker et al., 2021) and warm temperatures increased the abundance of ELF3 (Fig. 8B and D) (see also Ding *et al*., 2018; Zhang *et al*., 2021). This observation suggests that condensation in speckles protects ELF3 from degradation, as is the case of phyB in nuclear bodies (Rausenberger et al., 2010; Van Buskirk et al., 2014).

The formation of speckles by ELF3 itself showed hysteresis, as revealed by a significant shift in the sensitivity to temperature, when this is decreased as compared to when it is increased (Fig. 9D). Since the pattern of *PIF4* promoter activity requires ELF3 and correlates with ELF3 speckle formation (Fig. 9A, C, F), we conclude that ELF3 achieves hysteresis and drives the *PIF4* promoter into the same behaviour. What are the mechanisms that generate ELF3 hysteresis? Nuclear speckles are condensates, membrane-less compartments that may be formed by liquid-liquid phase separation (Emenecker et al., 2021). ELF3 undergoes phase separation to form nuclear speckles under warm temperatures; and both, *in vitro* phase separation and nuclear speckle formation depend on the intrinsically disordered prion-like domain of the protein (Jung et al., 2020). Hysteretic phase separation of intrinsically disordered proteins can emerge from intermolecular interactions that stabilise the aggregated phase (Quiroz et al., 2019); i.e., the origin of these hysteretic patterns could be at the ELF3 molecule itself. In fact, in response to increasing temperatures, purified ELF3 prion domain peptides form liquid droplets *in vitro*, and reversibility by temperature decreases is shifted towards lower temperatures (Jung et al., 2020), as observed here for *in vivo* speckle formation (Fig. 9D). Furthermore, ELF3 retained the memory of warm temperatures even before it formed speckles *in vivo* (Fig. S5). The so-called ‘mnemons’ are proteins that oligomerise to form condensates and establish long-lasting signalling changes, encoding a memory of previous conditions (Reichert and Caudron, 2021). There is abundant evidence for the function of these assemblies in yeast and Drosophila (Reichert and Caudron, 2021).

In conclusion, the history-dependent behaviour of PIF4 and HY5 and their upstream regulators ELF3 and COP1 enabled a memory of past temperature conditions. Some of these components, such as PIF4 and ELF3, drive this memory in one direction, as cooling barely affected the status established by warmth. Temperature is a variable aspect of the environment and here we show that cues provided by warm days and warm nights operate synergistically. Thus, integration of temperature information from different phases of the day has a main role in enabling plants attenuating their response to transitory oscillations in temperature (Fig. 1A, Table S1A).

## Materials and methods

### Plant material

We used *Arabidopsis thaliana*. The experiments where we measured hypocotyl growth or cotyledon expansion included the WT Columbia (Col-0). We list the mutant alleles and transgenic reporter lines in Table S2.

### Growth conditions

We used clear plastic boxes (4 × 3.5 × 2 cm^3^ height) for growth (14 seeds per genotype and box), microscopy (5 seeds per box) and protein blot (80 seeds per box) and microtiter plates for bioluminescence experiments (one seed per well). The substrate was 1.5 % agar-water. After sowing, we incubated the seeds for 4 days at 4 °C in darkness and transferred the stratified seeds to white light at 90 μmol m^-2^ s^-1^ (400-700 nm), provided by a mixture of fluorescent and halogen lamps, with a red/far-red ratio typical of sunlight (1.1), a photoperiod of 10 h (short day), and 20°C for 4 days. At the beginning of the fourth photoperiod (ZT= 0h), the seedlings received either white light or simulated shade (9 μmol m^-2^ s^-1^ between 400 and 700 nm with a red/far-red ratio of 0.1) at 10°C, 20°C or 28°C. For simulated shade, we combined the white light source with two green acetate filters (LEE filters 089). The seedlings initiated the night (ZT= 10 h) with or without 10 min of far-red light at 7 μmol m^-2^ s^-1^ (EOD FR), provided by 150 W incandescent lamps (R95, Philips) in combination with yellow, orange and red acetate filters (LEE filters 101, 105 and 106, respectively) and six blue acrylic filters (Paolini 2031, Buenos Aires, Argentina). Night temperature was 10°C, 20°C or 28°C. To investigate the occurrence of hysteresis without involving a light-to-dark transition, in some experiments, 4 h after the beginning of the fourth photoperiod (ZT= 4h), we introduced temperature shifts while the plants remained under white light.

### Hypocotyl growth rate

We photographed the seedlings with a digital camera (PowerShot; Canon, Tokyo, Japan) at the beginning of the night (ZT= 10 h) of the fourth photoperiod and either at the end of the night (ZT= 24 h) or at intermediate times (ZT= 14 h, 18 h and 22 h). We measured hypocotyl length increments by using an image processing software as described (Legris et al., 2016).

### Bioluminescence

We detected luciferase (LUC) activity with a Centro LB 960 (Berthold) luminometer by adding 20 μL of 0.2 mM D-luciferin per well 24 h before starting the measurements.

### Confocal microscopy

We obtained confocal fluorescence images from the epidermis and the first sub-epidermal layers of either the upper third portion of the hypocotyl (PIF4, HY5 and COP1) or individual nuclei present in the same region (ELF3) with a LSM5 Pascal laser-scanning microscope (Zeiss). The water-immersion objective lens were C-Apochromat X40/1.2 or C-Apochromatic X63/1.2 (Zeiss), respectively. We used an argon laser (λ= 488 nm) for excitation of GFP or YFP and a BP 505-530 filter for detection of fluorescence. We used a He-Ne laser (λ = 543 nm) for excitation of chlorophyll and a LP 560 filter for detection of its fluorescence and configured a transmitted light channel to visualise cellular structures. We performed image analysis in batch with an image segmentation program developed in Icy (http://icy.bioimageanalysis.org/) (Sellaro et al., 2019).

### Protein blots

We extracted total protein from 100 mg of seedlings by homogenizing plant material in extraction buffer containing 50 mM Tris-HCl (pH 7.5), 200 mM NaCl, 10 % (v/v) glycerol, 0.1 % (v/v) Tween-20, 1 mM PMSF and protease inhibitors (Roche), centrifuged twice (20 000 *g*, 15 min, at 4 °C), and transferred the supernatant into fresh Eppendorf reaction tubes. The protein concentration in the supernatant was determined by the Bradford assay (Bio-Rad). Protein samples were boiled (95 °C, 5 min) in TMx4 loading buffer. We run 20 µg of protein in 8% SDS-PAGE followed by wet blotting (100 mM Tris/Glycine, 10 % MeOH, 1.5 h). Homogeneous protein transfer to nitrocellulose membranes (Whatman) was confirmed by Ponceau red staining. Membranes were blocked (5 % milk powder, 0.1 % Tween-20 in TBS, 2 h) and incubated with primary antibody (anti-HA, 1:1000, Sigma) overnight, followed by incubation with anti-rabbit HRP-conjugated antibody (1:5000, Sigma) during 2 h. For 35S:PIF4-HA we used an anti-HA Peroxidase (Roche) antibody. We used ImageJ for quantification of the bands.

## Supplemental data

The following materials are available in the online version of this article

**Supplemental Figure S1**. Effects of daytime temperature on nighttime growth in multiple *pif* mutants.

**Supplemental Figure S2**. Daytime temperature affects nuclear fluorescence driven by the *pPIF4:PIF4-GFP* transgene in the cotyledons.

**Supplemental Figure S3**. Warm temperatures post-transcriptionally enhance PIF4 abundance.

**Supplemental Figure S4**. Temperature effects on the time course of *TOC1, LHY* and *CCA1* promoter activities.

**Supplemental Table S1**. Detailed statistical analysis of the data.

**Supplemental Table S2**. Mutant and transgenic lines used in this study.

## Acknowledgments

The authors are grateful to Dr Christian Fankhauser (University of Lausanne) for providing seed samples of the lines *pPIF7:PIF7-3HA-tPIF7* and *p35S:PIF4-HA*.

## Funding

This work was supported by the Argentinian *Agencia Nacional de Promoción Científica y Tecnológica* (grants PICT-2016-1459 and PICT-2018-01695 to JJC), *Universidad de Buenos Aires* (grant 20020170100505BA to JJC) and the Spanish *Ministerio de Ciencia e Innovación* (grant BIO2017-90056-R to SP).

### Conflict of interest statement

None declared

GM and JJC conceived and designed the experiments. SP and CN provided insightful suggestions. GM, CN and RS performed the experiments. SP provided new reporter lines. GM, CN, RS, SP and JJC analysed the data. GM and JJC wrote the paper with input from the other authors.

